# Proteome-wide quantitative RNA interactome capture (qRIC) identifies phosphorylation sites with regulatory potential in RBM20

**DOI:** 10.1101/2021.07.12.452044

**Authors:** Carlos Henrique Vieira-Vieira, Vita Dauksaite, Michael Gotthardt, Matthias Selbach

## Abstract

RNA-binding proteins (RBPs) are major regulators of gene expression at the post-transcriptional level. While many posttranslational modification sites in RBPs have been identified, little is known about how these modifications regulate RBP function. Here, we developed quantitative RNA-interactome capture (qRIC) to quantify the fraction of cellular RBPs pulled down with polyadenylated mRNAs. Applying qRIC to HEK293T cells quantified pull-down efficiencies of over 300 RBPs. Combining qRIC with phosphoproteomics allowed us to systematically compare pull-down efficiencies of phosphorylated and non-phosphorylated forms of RBPs. Over hundred phosphorylation events increased or decreased pull-down efficiency compared to the unmodified RBPs and thus have regulatory potential. Our data captures known regulatory phosphorylation sites in ELAVL1, SF3B1 and UPF1 and identifies new potentially regulatory sites. Follow-up experiments on the cardiac splicing regulator RBM20 revealed that multiple phosphorylation sites in the C-terminal disordered region affect nucleo-cytoplasmic localization, association with cytosolic RNA granules and alternative splicing. Together, we show that qRIC is a scalable method to identify functional posttranslational modification sites in RBPs.

**Highlights:**

- qRIC globally quantifies the fraction of RNA-binding proteins pulled down with mRNA
- Combining qRIC with phosphoproteomics identifies sites that affect RNA binding
- Phosphorylation sites in RBM20 regulate its function in splicing

## Introduction

Cellular RNA-binding Proteins (RBPs) regulate RNA fate in a time- and space-dependent manner and thus play a key role in posttranscriptional gene expression control in health and disease (Gerstberger, Hafner and Tuschl, 2014; Wurth and Gebauer, 2015; Gehring, Wahle and Fischer, 2017; Hentze *et al*., 2018; Buccitelli and Selbach, 2020). The human genome is estimated to encode over 1,900 RBPs (Hentze *et al*., 2018), a number higher than the about 1,600 transcription factors (Lambert *et al*., 2018). While the activity of transcription factors is well-known to be regulated by cellular signal transduction cascades, we know relatively little about how cell signaling affects the function of mRNA-Binding Proteins (mRBPs).

Cell signaling often involves post-translational protein modifications (PTMs) - reversible, covalent additions like phosphorylation, acetylation or ubiquitination (Bludau and Aebersold, 2020). The most widely studied PTM is phosphorylation, and biochemical enrichment strategies combined with quantitative mass spectrometry-based proteomics can now identify thousands of phosphorylation sites in mammalian samples and more than 50,000 sites in a single human cell line (Dephoure *et al*., 2008; Macek, Mann and Olsen, 2009; Choudhary and Mann, 2010; Huttlin *et al*., 2010; Sharma *et al*., 2014; Bekker-Jensen *et al*., 2020). PTMs, particularly phosphorylation, are also known to regulate the activity of RBPs (Yu, 2011; Thapar, 2015; Lovci, Bengtson and Massirer, 2016; Zarnack *et al*., 2020). For example, phosphorylation of RBPs can affect translation (Imami *et al*., 2018; Jia *et al*., 2021), RNA stability and processing (Zhang *et al*., 2018; Kim and Maquat, 2019), RBP subcellular localization (Hyeon Ho Kim *et al*., 2008), and splicing (Stamm, 2008; Murayama *et al*., 2018). Also, PTMs play a role in liquid-liquid phase separation of ribonucleoprotein granules (Hofweber and Dormann, 2019; Nosella and Forman-Kay, 2021). Despite these insights, the functional relevance of most PTM sites for RBP function is unknown. More broadly speaking, while over 200,000 identified phosphorylation sites are reported in databases such as PhosphositePlus, the vast majority of them lack functional annotation (Hornbeck *et al*., 2015; Needham *et al*., 2019; Ochoa *et al*., 2020). This is mainly due to the fact that the experimental techniques used to assess the function of individual phosphorylation sites are typically not scalable. More recently, systematic approaches to identify functionally relevant PTM sites are beginning to emerge (Imami *et al*., 2018; Huang *et al*., 2019; Masuda *et al*., 2020; Smith *et al*., 2021).

The defining feature of RBPs is their ability to bind RNA. UV crosslinking can “freeze” such interactions *in situ* for subsequent protein- or RNA-centric analyses (Vieira-Vieira and Selbach, 2021). Protein-centric methods such as cross-linking and immunoprecipitation (CLIP) involve purification of an individual RBP and identification of its RNA binding sites by sequencing (Wheeler et al., 2018; Lin and Miles, 2019). RNA-centric methods like RNA-interactome capture (RIC) are based on biochemical isolation of RNAs followed by identification of the RNA-bound proteome by mass spectrometry (Smith et al, 2020; Gräwe *et al*., 2020). CLIP and RIC have provided us with useful catalogs of RBPs and their RNA-binding sites (Gerstberger, Hafner and Tuschl, 2014; Maatz *et al*., 2014; Hentze *et al*., 2018; Zhu *et al*., 2019; Van Nostrand *et al*., 2020). Also, comparative RIC studies provided information about how the interaction of RBPs with RNA changes upon perturbation (Milek *et al*., 2017; Perez-Perri *et al*., 2018; Garcia-Moreno *et al*., 2019; Hiller *et al*., 2020). However, the impact of PTMs in RBPs on their interaction with RNAs has not yet been investigated systematically.

We reasoned that combining RIC with quantitative mass spectrometry should enable us to quantify the pull-down efficiency of mRNA binding proteins (mRBPs) on a proteome-wide scale. Here, we develop quantitative RIC (qRIC) to quantify the fraction of cellular mRBPs pool that can be cross-linked to and co-purified with mRNA. Combined with phosphoproteomics, qRIC allowed us to assess how the modification state of individual sites correlates with its pull-down efficiency. We identify over hundred phosphorylation sites with regulatory potential in HEK cells, including both sites known to regulate mRBP function and novel candidate sites. Follow-up experiments on the cardiac splicing regulator RBM20 revealed that several phosphorylation sites in its C-terminal disordered region affect nucleo-cytoplasmic localization, association with cytosolic RNA granules and impact splicing function. In summary, we establish qRIC as a scalable method to assess the function of posttranslational modification sites in RBPs, apply it to assess phosphorylation in HEK293 cells and provide new insights into the function of RBM20.

## Results

### qRIC quantifies pull-down efficiency of mRBPs in association with mRNA in vivo

The mRNA-bound proteome can be characterized by RIC, i.e. photocrosslinking of mRBPs to mRNAs in living cells, followed by oligo(dT) purification (Hentze *et al*., 2018; Gräwe *et al*., 2020). Covalent cross-linking enables high-stringency washes that remove essentially all contaminating proteins (Baltz *et al*., 2012; Castello *et al*., 2012). Hence, while other affinity purification experiments often require quantification to distinguish genuine interaction partners from non-specific contaminants (Meyer and Selbach, 2015; Smits and Vermeulen, 2016), RIC experiments do not necessarily depend on quantification as an additional filter. Instead, comparative RIC (cRIC) experiments use relative quantification to compare the RNA-bound proteome in different conditions in order to assess RNA-binding changes upon perturbation (Milek *et al*., 2017; Perez-Perri *et al*., 2018; Garcia-Moreno *et al*., 2019; Hiller *et al*., 2020). We reasoned that quantifying the fraction of proteins co-purifying with mRNAs relative to their overall cellular abundance in cells would be particularly informative, especially to assess the impact of phosphorylation. For example, if phosphorylation of a specific site in an mRBP affects mRNA binding, the fraction of the phosphorylated protein that is pulled-down via RIC should differ from the fraction of the non-modified protein.

Based on this idea, we designed quantitative RIC (qRIC) to systematically assess mRBP pull-down efficiencies (Fig.1 A). In qRIC, cells are first differentially labelled with light of heavy stable isotopes using SILAC (Mann, 2006). Protein-RNA interactions are then stabilized *in vivo* by UV cross-linking. From the heavy labelled cells, the RBPs which are covalently bound to mRNAs are pulled down with magnetic oligo(dT) beads followed by stringent washes. These heavy labeled samples (“Pull-down”) are combined with 1% of the whole cell protein extract (“Input”) from light cells as an internal reference to enable accurate quantification. After protein digestion, peptide mixtures are analyzed by quantitative shotgun proteomics. The fraction of the total cellular protein pool that is pulled down with oligo(dT) beads (that is, the pull-down efficiency) is determined from the SILAC ratios relative to the light spike-in input sample. In this way, qRIC directly associates heavy to light SILAC ratios with the pull-down efficiencies. In parallel to whole proteome samples, a fraction of the peptide mixtures is used to enrich phosphorylated peptides by TiO2 chromatography, followed by shotgun proteomics. Comparing the pull-down efficiencies of individual phosphorylated peptides with the efficiencies of the corresponding mRBPs (that is, calculating the delta efficiency - ratio of SILAC ratios) then identifies phosphorylation sites with regulatory potential.

**Figure 1.**
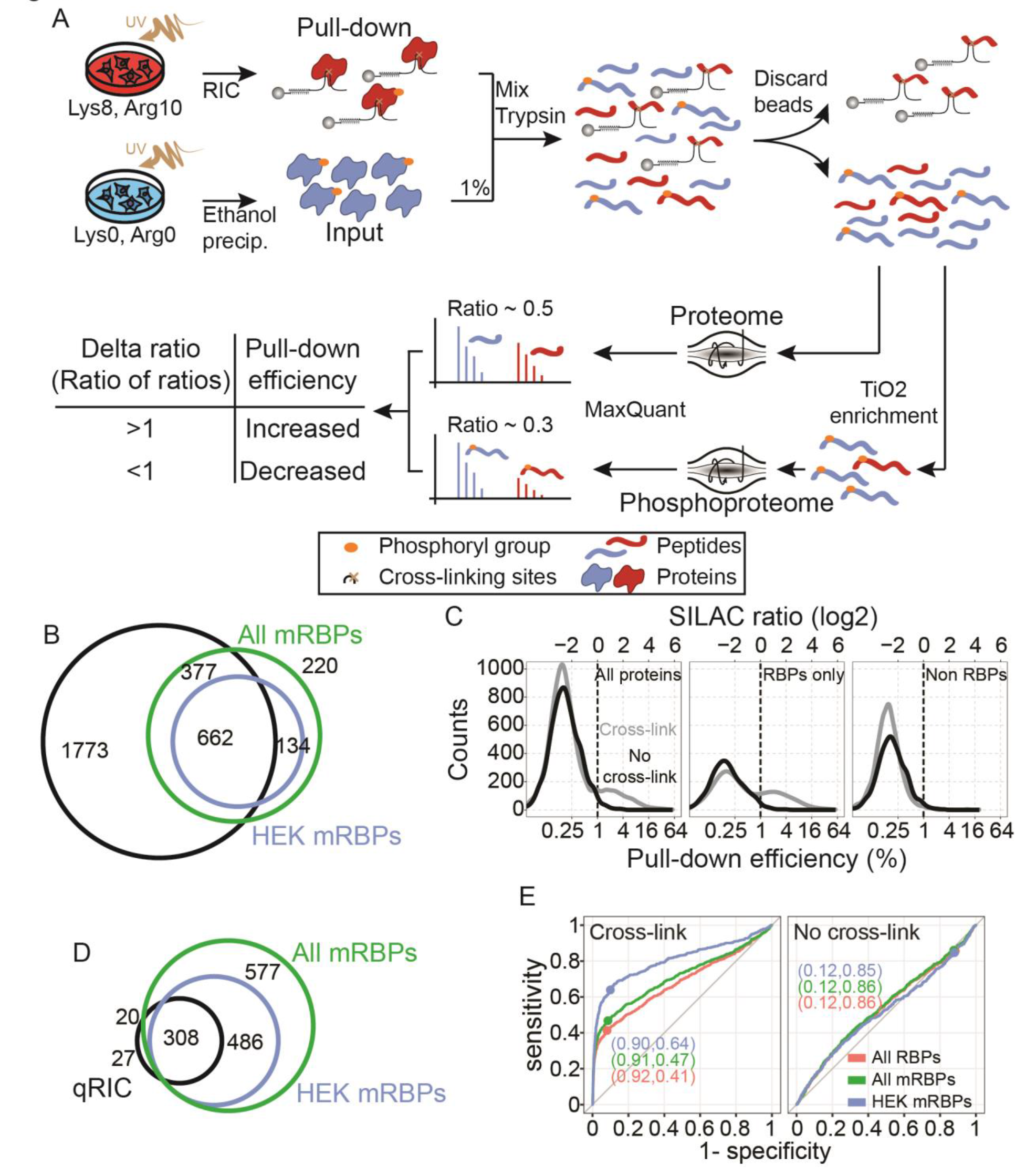
Quantification of mRBPs pull-down efficiencies with qRIC. (A) Experimental design for qRIC. See also Figure S1. (B) Venn diagrams of proteins identified in our dataset (black), HEK-specific mRBPs (green) and all annotated mRBPs (blue). (C) Proteins pull-down efficiencies from UV cross-linked (grey) and non-crosslinked cells (black). Annotated RBPs and non-RBPs are re-plotted to highlight the RBP-specific pull-down efficiency peak. Dashed lines indicate the 1 % pull-down efficiency threshold. (D) Same as B but only proteins quantified with pull-down efficiency higher than 1 % in both forward and reverse biological replicates were selected. (E) Receiver operating characteristic curves for RBP calling using different sets of RBPs. Values for specificity and sensitivity at the threshold of 1 % pull-down efficiency are indicated in parentheses.

To establish qRIC we initially included an additional medium-heavy control without UV cross-linking (Fig. S1A). This experiment was carried out in biological duplicates with swapped isotope labels. As expected, the well-known mRBP HNRNPD was specifically pulled-down in the UV-irradiated sample but not in the non-crosslinked control (Fig. S1B, C). Spiking in 1% of the total input (“L” in Fig. S1B and D, and “H” in S1C and E) revealed a pull-down efficiency for HNRNPD of 6%. In contrast, a protein like Actin (ACTN1) was mainly detected in the input with only background signal in the UV-irradiated and non-UV-irradiated samples and therefore had low pull-down efficiency (Fig. S1D, E). Pull-down efficiencies in the forward and reverse label experiments were correlated, indicating overall good biological reproducibility (Fig. S1 F).

Encouraged by these results, we performed four additional qRIC experiments as outlined in Fig. 1 A (that is, omitting the non UV-crosslinked control). Again, label swap experiments were highly correlated (Fig. S1G). In total, we identified 5,461 proteins in all six biological replicates combined. We required proteins to be quantified in at least one of the three forward and one of the three reverse experiments to be considered for further analysis. We computed the mean of all three forward and all three reverse experiments, followed by taking the mean-of-means from forward and reverse experiments to obtain a single average pull-down efficiency for 2831 proteins. The entire dataset is available as supplementary table 1 (Table S1).

To assess coverage we compared our data to previously identified RBPs (Baltz *et al*., 2012; Hentze *et al*., 2018). We covered 75% (1039 out of 1393) and 83% (662 out of 796) of all mRBPs and of cell line-specific mRBPs, respectively (Fig. 1 B). Plotting mean pull-down efficiencies for all 2831 quantified proteins showed a bimodal distribution with one peak at about 0.2 and another one at 2% (Fig. 1 C). The peak at 0.2% was essentially unaltered in the control experiments without UV cross-linking while the peak at 2% was markedly reduced. Hence, the low efficiency peak appears to be largely due to non-specific contaminants (that is, proteins co-purifying with oligo(dT) beads that are not covalently cross-linked to mRNAs). Consistently, the vast majority of proteins with pull-down efficiencies greater than 1% are annotated as RBPs (Fig. 1 D) (Hentze *et al*., 2018). Systematic benchmarking using annotated RBPs as a reference revealed that qRIC identifies mRBPs with good sensitivity and specificity (Fig. 1 E). For subsequent analyses, we chose a conservative threshold of 1% pull-down efficiency (in both forward and reverse experiments) to ensure high specificity (90%).

### Features affecting mRBP pull-down efficiencies

The pull-down efficiency depends on factors such as UV crosslinking and oligo(dT) enrichment and should therefore not be interpreted as a direct readout for the actual fraction of an RBP bound to mRNAs *in vivo* (Vieira-Vieira and Selbach, 2021). Bearing this limitation in mind, we asked if pull-down efficiencies correlate with specific RBP features. For proteins that passed our threshold we observed a small but significant positive correlation of pull-down efficiencies with absolute cellular protein copy number estimates (Fig. 2 A) and negative correlation with protein size (Fig. 2 B). To assess how the presence of annotated RNA-binding domains (RBDs) correlates with pull-down efficiencies we extracted domain annotation from the Pfam database (Mistry *et al*., 2021a) and selected domains annotated as RNA-binding (Gerstberger, Hafner and Tuschl, 2014). We found that the number of RBDs per protein strongly impacts pull-down efficiency (Fig. 2 C). Pull-down efficiencies of proteins with and without annotated RBDs differed already below the 1% threshold, confirming that our cut-off is indeed conservative. For all proteins containing only a single RBD we also compared pull-down efficiencies for different RBDs. Proteins with single KH_1 and RRM_1 domains showed higher pull-down efficiencies than proteins with a single DEAD box domain, consistent with the more transient nature of DEAD box helicases binding to RNA (Fig. 2 D).

**Figure 2.**
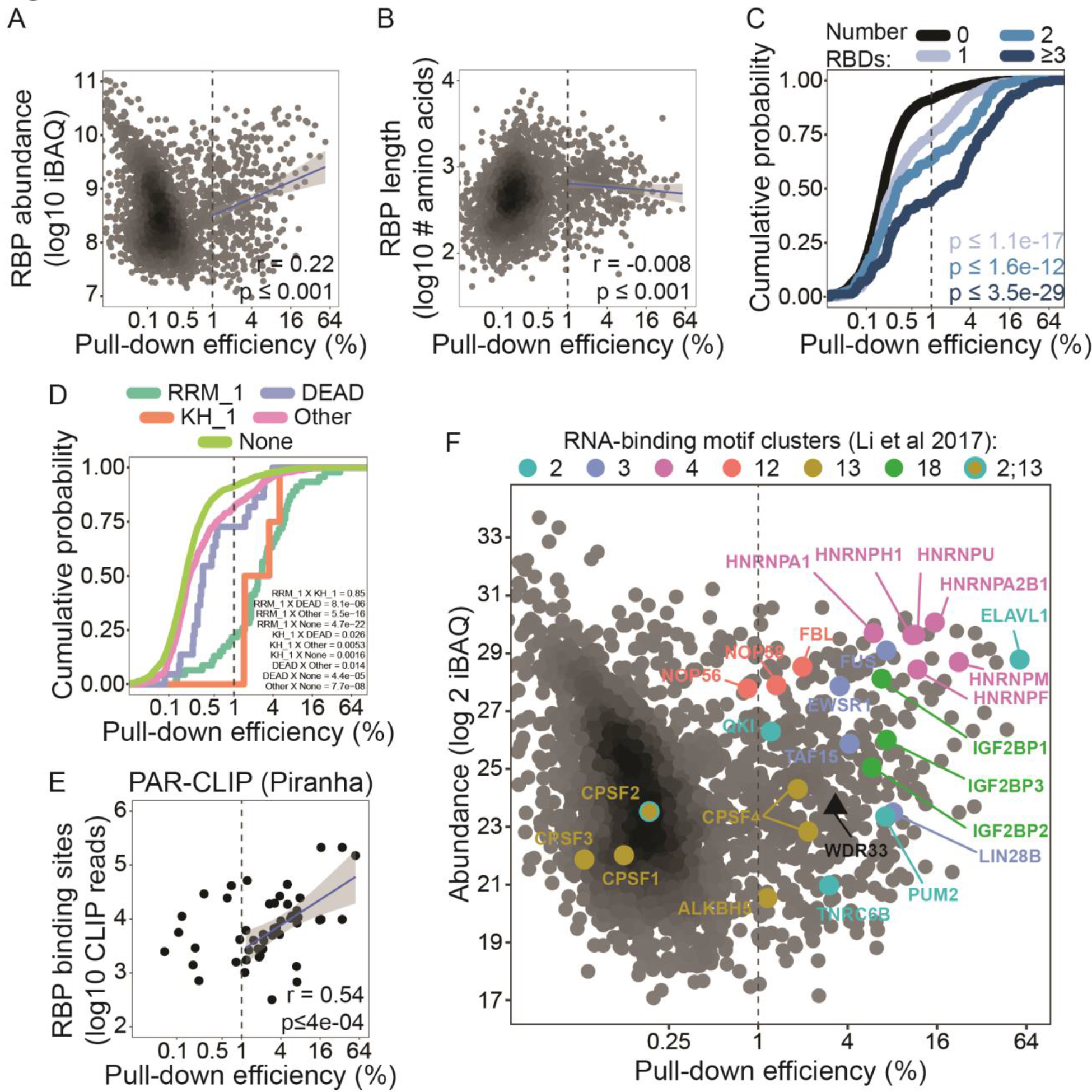
Features affecting mRBP pull-down efficiencies. (A and B) Protein pull-down efficiency is plotted against proteome absolute abundance (iBAQ intensity) and protein length (number of amino acids), respectively. Pearson correlation was performed for proteins with pull-down efficiency higher than 1 % in both forward and reverse biological replicates. R correlation value is indicated. P-value was estimated from ten thousand randomized assignments of the data. (C and D) Pull-down efficiency cumulative probability of proteins grouped according to the number and type of annotated RBDs. In C, p-values from a two-sided Wilcoxon test for the comparison with proteins without annotated RBDs is indicated. In D, only proteins with a single RBD are plotted and p-values from a two-sided Wilcoxon test are indicated for each group comparison. (E) Number of peaks from PAR-CLIP data produced in HEK293 cells and analysed with the Piranha peak calling tool. Pearson correlation was performed for proteins with pull-down efficiency higher than 1 % in both forward and reverse biological replicates. R correlation value is indicated. P-value was estimated from ten thousand randomized assignments of the data. See also Figure S2. (F) Same as A but proteins belonging to RNA-binding motifs are highlighted. WDR33 is emphasized (see main text).

To compare our RNA-centric (that is, RIC) with protein-centric (that is, CLIP) data we took advantage of a database for CLIP-seq experiments (Zhu *et al*., 2019). Here, we analysed the total number of peaks in PAR-CLIP data produced in HEK293 and HEK293T cells as a proxy for the number of RNA-binding sites in the transcriptome. We found a clear correlation between protein pull-down efficiency and the total number of CLIP-seq peaks (Fig. 2 E). This finding is independent of the peak calling tool employed (Fig. S2 A). Interestingly, the correlation was lower for eCLIP data obtained in K562 cells and HepG2 cells, indicating that pull-down efficiencies are to some extent cell type dependent (Fig. S2 B and C).

Different RBPs can bind to similar RNA motifs. We therefore compared pull-down efficiencies of RBPs grouped together based on their shared binding sites via combinatorial clustering (Li *et al*., 2017). Interestingly, we find that RBPs interacting with similar sequence motifs tend to have similar pull-down efficiencies, highlighting the importance of the number of RNA-binding sites for pull-down efficiencies (Fig. 2 F). This is even true for proteins with vastly different absolute abundance in HEK293T cells such as LIN28B and FUS. In contrast, CPSF4 was more efficiently pulled down than CPSF1, CPSF2 and CPSF3, even though all four proteins are annotated to interact with the same motif (Li *et al*., 2017). All four proteins are members of CPSF (cleavage and polyadenylation specificity factor), a multiprotein complex that recognizes the AAUAAA motif in the polyadenylation signals of precursor mRNAs (Keller *et al*., 1991). While initially CPSF1 was thought to directly bind to this sequence motif, more recent data showed that RNA-binding occurs via CPSF4 and WDR33 instead (Keller *et al*., 1991; Chan *et al*., 2014; Schönemann *et al*., 2014; Clerici *et al*., 2018). This is consistent with our observation that CPSF4 but none of the other CPSF proteins is efficiently pulled down. In addition, WDR33 is pulled down with a similar efficiency to CPSF4 (Fig. 2 F). These observations highlight the strength of qRIC to distinguish direct from indirect binders. In summary, we conclude that pull-down efficiencies determined via qRIC provide a meaningful read-out for RNA-protein interaction.

### Quantifying the impact of phosphorylation on mRBP pull-down efficiency in qRIC

We next focused our attention on the analysis of phosphorylation sites. Similar to proteins, pull-down efficiencies of phosphorylated peptides identified in qRIC also showed a bimodal distribution (Fig. 3 A). The higher efficiency peak disappeared in the control group without UV cross-linking and mainly consists of phosphorylated peptides from annotated RBPs. Hence, also the phosphorylated peptide data reflects direct UV cross-linking dependent RBP-RNA interactions. In all six biological replicates combined, 2719 class I (localization probability > 0.75) phosphorylation sites were identified and pull-down efficiencies for 1243 were quantified, mostly on serine residues (Fig. 3 B, Supplementary Table S2). In total, 479 phosphorylation sites on 196 proteins had pull-down efficiencies >1% in both forward and reverse SILAC label experiments. For subsequent analyses, we only considered reproducibly quantified phosphorylation sites for which the corresponding host protein was also reproducibly quantified and either the phosphorylated peptide or the host protein passed the 1% pull-down efficiency threshold (see Methods). This yielded 395 phosphorylation sites in 166 RBPs (Fig. 3 B).

**Figure 3.**
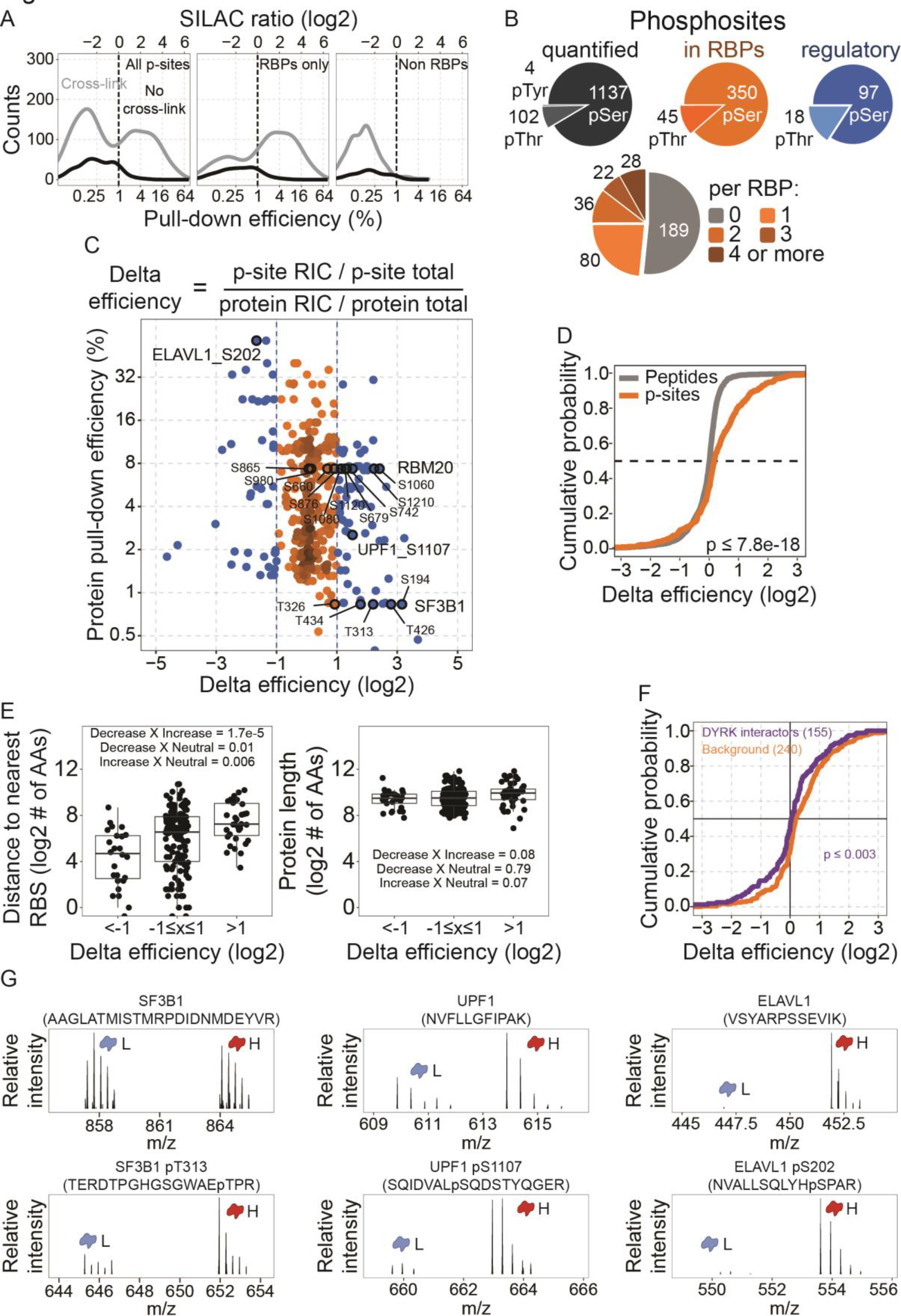
Quantifying the impact of phosphorylation on mRBP pull-down efficiency in qRIC. (A) Same as Figure 1 C but phosphorylation sites (p-sites) are shown instead of proteins. (B) Overview of quantified phosphorylation sites in qRIC. (C) Phosphorylation site delta efficiency plotted against the pull-down efficiency of host proteins. Candidate regulatory phosphorylation sites are colored in blue. Phosphorylation sites discussed in the main text are highlighted. The delta efficiency formula is also shown. (D) Delta efficiency cumulative probability of phosphorylated peptides and unmodified peptides from host proteins. Only unmodified peptides from proteins with at least one phosphorylation site were considered. The p-value from a two-sided Wilcoxon test for the comparison between groups is indicated. (E) Phosphorylation site linear distance to the nearest RNA-binding site (RBS) and length of host proteins in number of amino acids. Groups were compared using one-way ANOVA and Tukey’s post-test p-values are shown for each comparison. (F) Delta efficiency cumulative probability of phosphorylation sites in DYRK3 targets. The p-value from a two-sided Wilcoxon test for the comparison to the group of phosphorylated sites in non-DYRK3 targets is indicated. The number of phosphorylation sites in each group is also indicated in parentheses. (G) The phosphorylated peptide spectra (bottom row) and exemplary unmodified peptide spectra (top row) for SF3B1 T313, UPF1 S1107 and ELAVL1 S202 phosphorylation sites. SILAC light (L) and heavy (H) labeled chromatogram peaks are indicated. The peptide sequences are also indicated in parentheses.

To systematically identify phosphorylation sites with regulatory potential, we compared the pull-down efficiencies of individual sites with the efficiency of the corresponding host protein. To this end, we computed delta efficiencies as the ratio of the H/L SILAC ratios (Fig. 3 C, Supplementary Table S3): Positive and negative log delta efficiencies indicate that phosphorylation of the respective site correlates with increased or decreased pull-down efficiency, respectively. In total, 73 and 42 phosphorylation sites correlate with at least 2-fold increased and decreased pull-down efficiency, respectively. The cumulative distribution of delta efficiencies for phosphorylated and non-phosphorylated peptides revealed that phosphorylated peptides tend to show increased mRNA interaction in qRIC (Fig. 3 D). This is surprising since phosphorylation adds negative charges to proteins and might thus be expected to inhibit binding of RBPs to the negatively charged RNA. To assess the impact of phosphorylation sites that are located close to the actual RNA-protein interaction interface, we took advantage of a recently published study that used chemical RNA digestion to identify the RBP residues cross-linking to RNA (Bae *et al*., 2020). Here, we observed that sites with negative delta efficiencies tended to be closer to known cross-linking residues (in terms of primary sequence length), although the effect was small (Fig. 3 E). This is consistent with the idea that phosphorylation sites that are close to RNA-binding regions tend to be more inhibitory. However, we note that our analysis is biased against detection of activating phosphorylation sites near cross-linking residues since the peptides within an RBP that are covalently cross-linked to the RNA are lost in our qRIC workflow.

Phosphorylation and other PTMs are also known to be involved in liquid-liquid phase separation of membraneless organelles such as various types of RNA granules (Owen and Shewmaker, 2019; Nosella and Forman-Kay, 2021). For example, the kinase DYRK3 acts as a central dissolvase of several types of membraneless organelle like stress granules and splicing speckles during mitosis (Rai *et al*., 2018). Many DYRK3 interaction partners are RNA-binding proteins and members of such ribonucleoprotein particles. Interestingly, we found that phosphorylation sites in DYRK3 interaction partners had mildly but significantly reduced delta efficiencies (Fig. 3 F). This is consistent with the idea that DYRK3 dissolves membraneless organelles by phosphorylating RBPs, thus inhibiting their interaction with RNAs.

### qRIC data reflects known regulatory phosphorylation sites

Apart from the global trends outlined above, the main strength of the data is that it provides functional information for individual phosphorylation sites. Indeed, taking a closer look at the qRIC data revealed that it reflects the known function of several phosphorylation sites.

First, we observed that five phosphorylation sites in SF3B1 correlate with increased mRNA binding in qRIC. SF3B1 (a.k.a. SF3B155 or SAP155) is a member of the U2 splicing complex and directly binds mRNA in activated spliceosomes (Cretu *et al*., 2016). Several SF3B1 phosphorylation sites regulate splicing, and phosphorylation of one of the sites we identified (T313) is an established marker for spliceosome activity (Wang *et al*., 1998; Boudrez *et al*., 2002; Girard *et al*., 2012). The pull-down efficiency of pT313 was ∼6% compared to only ∼1% of the SF3B1 protein (Fig. 3 G) yielding a delta efficiency of 4.6. The other four sites quantified in qRIC are all located in the same intrinsically disordered N-terminal domain of SF3B1 that is not part of the core region of the SF3b complex and thought to have a regulatory function during the splicing cycle (Cretu et al. 2016).

Second, we found that phosphorylation of S1107 in UPF1 correlates with increased mRNA binding (Fig. 3 G). UPF1 is an RNA-dependent helicase and ATPase playing a central role in the nonsense-mediated mRNA decay (NMD) and other mRNA degradation pathways (Kurosaki, Popp and Maquat, 2019). Importantly, phosphorylation of UPF1 by its kinase SMG-1 is known to require association with target mRNAs (Kashima et al., 2006). Phosphorylated UPF1 recruits the CCR4−NOT complex to initiate mRNA degradation, and inhibition of mRNA degradation leads to accumulation of S1107-phosphorylated UPF1 (Durand, Franks and Lykke-Andersen, 2016). Thus, our data reflects the known role of this phosphorylation site for UPF1 function.

Third, we found that phosphorylation of S202 in ELAVL1 correlated with decreased mRNA binding (Fig. 3 G). ELAVL1 (a.k.a. HuR) is a well-characterized RBP that binds AU-enriched elements in the 3′-UTR to control splicing, localization, stability, and translation of target mRNAs (Grammatikakis, Abdelmohsen and Gorospe, 2017). ELAVL1 function is affected by posttranslational modifications that regulate its subcellular localisation. Upon phosphorylation on S202, ELAVL1 interacts with nuclear 14-3-3 proteins and is retained in the nucleus, reducing its association with cytosolic mRNA targets (H. H. Kim *et al*., 2008). Hence, the reduced association of ELAVL1 phosphorylated on S202 is consistent with the known regulatory function of this phosphorylation site. In conclusion, these examples highlight that qRIC can identify regulatory phosphorylation sites involved in diverse post-transcriptional gene regulatory events.

### qRIC identifies regulatory phosphorylation sites in RBM20

We next focused our attention on sites that have not yet been functionally characterized. Intriguingly, six phosphorylation sites in RBM20 (S679, S742, S1060, S1080, S1120 and S1210) correlated with significantly increased pull-down efficiency. Two additional sites (S660 and S876) showed the same tendency but did not pass the cut-off (Fig. 3 C), while another two sites showed no difference (S865 and S980).

RBM20 is a vertebrate-specific RBP with a key role in regulating alternative splicing (Watanabe, Kimura and Kuroyanagi, 2018; Lennermann, Backs and van den Hoogenhof, 2020). First insights into its function came from the observation that mutations in RBM20 are linked to familial forms of dilated cardiomyopathy (DCM) (Brauch *et al*., 2009). Follow-up studies revealed that RBM20 regulates alternative splicing of transcripts encoding key structural proteins in muscle cells, including the giant protein titin (Guo *et al*., 2012). Wild-type RBM20 leads to exclusion of specific exons in target transcripts, and pathogenic mutations in RBM20 as well as low expression levels cause erroneous exon retention (Guo *et al*., 2012). The protein contains two zinc finger (ZnF) domains, an RNA-recognition motif (RRM)-type RNA-binding domain and an arginine/serine (RS)-rich region with most pathogenic mutations in an RSRSP stretch in the RS region (Lennermann, Backs and van den Hoogenhof, 2020). The two serine residues in this stretch are phosphorylated in rodents and required for RBM20 nuclear localisation (Murayama *et al*., 2018; Sun *et al*., 2020).

The ten phosphorylation sites mentioned above are found in the disordered C-terminal half of the protein outside the RS region (Fig. 4 A). This region is essential for RBM20 mediated splicing repression (Dauksaite and Gotthardt, 2018) but not for the nuclear localization of the protein (Filippello *et al*., 2013). To characterize the role of these phosphorylation sites we created phosphomimetic and non-phosphorylatable mutants by exchanging all ten serine to aspartic acid (D10 mutant) and alanine (A10 mutant) residues, respectively. For comparison, we also used the well-characterized disease-causing S635A mutant in the RS domain that fails to regulate splicing (Guo *et al*., 2012; Maatz *et al*., 2014; Murayama *et al*., 2018).

**Figure 4.**
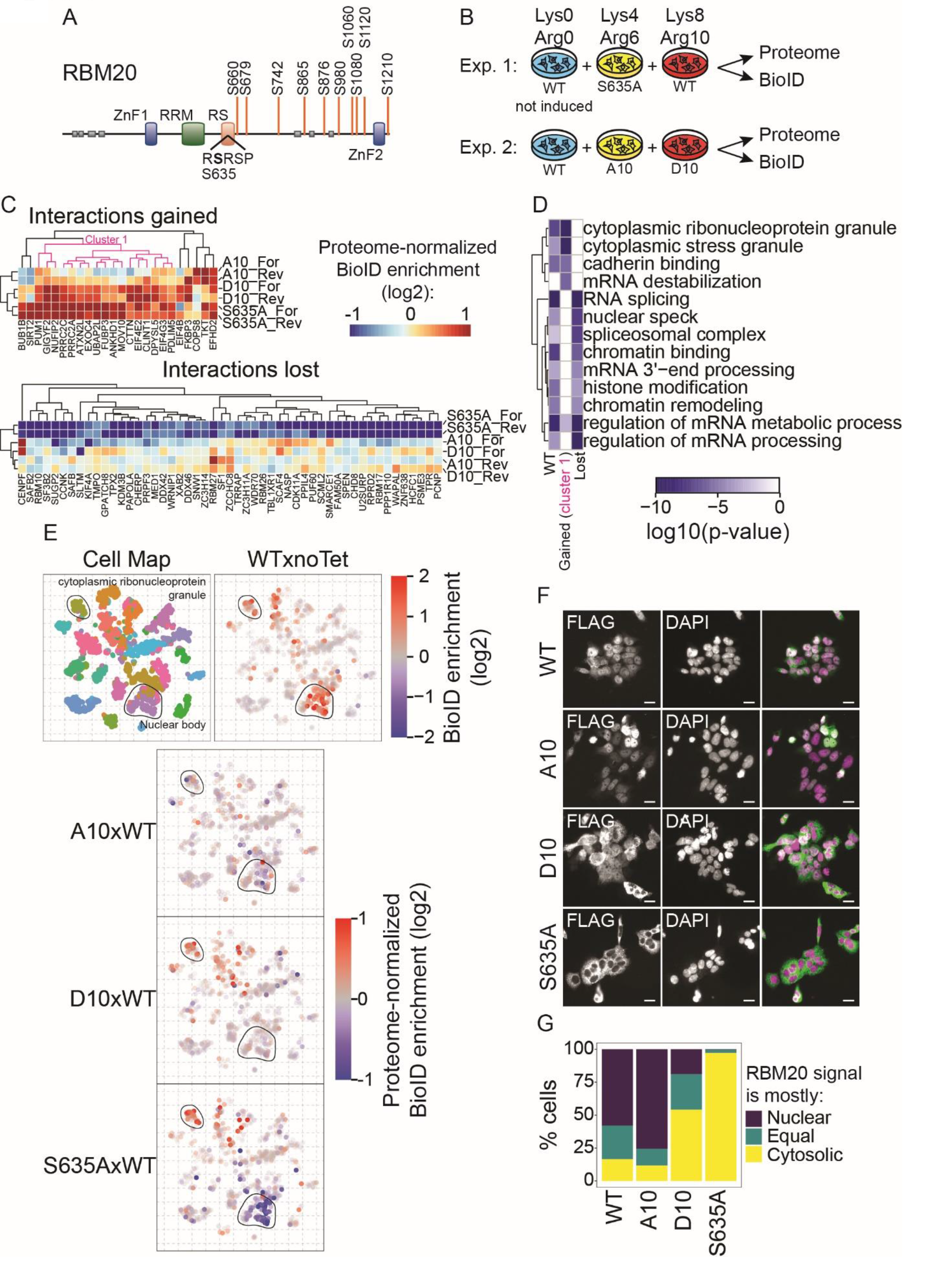
BioID investigation of candidate regulatory phosphorylation sites in RBM20. (A) Schematic representation of the human RBM20 protein and regulatory phosphorylation sites (see Figure 3). A central black line represents protein sequence with colored boxes indicating annotated protein domains. Gray boxes indicate disordered regions. Vertical orange bars indicate the position of regulatory phosphorylation sites. (B) Design of BioID experiments. Protein SILAC ratios in total and BioID enriched proteomes were quantified by mass spectrometry. See also Figure S3. (C) Hierarchical clustering for RBM20 gained and lost interactions in mutant variants compared to wild-type. Mutant-over-wild-type RBM20 BioID enrichment ratios were normalized by proteome changes. Cluster 1 denotes gained interactions by both D10 and S635A but not A10. (D) Hierarchical clustering for GO terms enriched in wild-type RBM20 proximity interactors (WT) and interactions gained or lost by mutants. (E) Projection of BioID results onto the tSNE plot proximity map. The original proximity map is shown and compartments of interest highlighted for reference. (F) Immunofluorescence microscopy imaging in cells expressing the FLAG-tagged, BioID-fused RBM20 variants. RBM20 was detected by immunostaining against the FLAG tag. DAPI was used to stain the nucleus. (G) Quantification of results in F.

To systematically compare the cellular neighborhood of different RBM20 mutants we used BioID, a proximity-dependent biotinylation method (Roux *et al*., 2012; Gingras, Abe and Raught, 2019). We created stable HEK293 cell lines expressing RBM20 mutants fused to a BirA-FLAG tag under the control of a tetracycline inducible promoter (Couzens *et al*., 2013). We then quantified differences in proximity-dependent biotinylation across conditions in biological duplicates (Fig. 4 B). Comparing cells expressing BirA-FLAG wild-type (WT) RBM20 with non-induced controls without tetracycline revealed proximity-dependent biotinylation of 95 proteins, including several splicing related factors like SF1, SF3B2, SAFB, SAFB2, U2SURP, DDX46, RBM10 and RBM17 (Fig. S3 A). Gene ontology analysis confirmed specific enrichment of terms like “mRNA processing” and “chromatin remodeling”, consistent with the nuclear localisation of WT RBM20 and its function in splicing (Fig. 4 D). We then asked how RBM20 mutations affect proximity-dependent biotinylation relative to the wild-type. To avoid biases caused by differences in steady-state protein levels between cell lines, we also quantified whole proteomes and only considered proteins with proteome-normalized, proximity-dependent biotinylation changes (Fig. S3 B). The pathogenic S635A mutant showed markedly reduced interactions with splicing factors, consistent with its impaired splicing regulatory function (Fig. 4 C). In contrast, the D10 and A10 mutations had little impact on RMB20 interaction with splicing-related proteins. Conversely, both the S635A and D10 but not the A10 mutant showed increased interactions with proteins enriched in the GO term “cytoplasmic stress granule” (Fig. 4 D).

To further characterize our BioID data we took advantage of a recently published protein proximity map of HEK293 cells (Go *et al*., 2021). Projecting our WT RBM20 data onto this map revealed enrichment signals at different subcellular locations (Fig. 4 E). When we mapped the changes of the S635A mutant relative to the wild-type onto the map we observed a decrease in “nuclear body” and an increase in “cytoplasmic ribonucleoprotein granule” signal. This is consistent with the decreased interaction of this mutant with splicing factors and its increased localisation to cytosolic granules (Maatz *et al*., 2014; Sun *et al*., 2020). In contrast to the S635A mutant, the A10 and D10 mutants showed only minor changes in “nuclear body”. Importantly, however, the D10 mutant displayed an increased signal in the “cytoplasmic ribonucleoprotein granule” compartment.

In summary, the GO- (Fig. 4 D) and the proximity map-based (Fig. 4 E) analyses consistently indicate that the phosphomimetic D10 mutant and the pathogenic S635A mutant associate increasingly with cytosolic stress granules. While the S635A mutant also shows reduced interaction with nuclear splicing-related proteins, this is not the case for the D10 and A10 mutants. The behaviour of the A10 mutant was overall very similar to wild-type RBM20. To validate these observations we also performed immunofluorescence microscopy (Fig. 4 F and G). Consistent with the BioID data, both the wild-type and the A10 mutant were mostly found in the nucleus. The S635A mutant was depleted in the nucleus and almost exclusively cytosolic. While the D10 was also more cytosolic than the wild-type, a considerable fraction of the protein was still localized to the nucleus. Hence, phosphorylation of RBM20 appears to cause a partial relocation of the protein to cytosol, reminiscent of the pathogenic S635A mutant.

### Phosphomimetic RBM20 mutations impair its splicing regulatory function

Since the pathogenic S635A mutant fails to regulate splicing of target transcripts, we asked how the phosphomimetic and non-phosphorylatable mutations affect splicing. To this end, we used two cell-based splicing reporter systems. The first system is based on a transiently transfected titin construct containing sequences encoding firefly and *Renilla* luciferases (Fluc and Rluc, respectively) inserted in exons 8 and 13 of the titin PEVK region (Fig. 5 A). In this system, wild-type RBM20 leads to the exclusion of the Fluc-containing exon and a decreased Fluc to Rluc ratio (Guo *et al*., 2012). While the A10 mutant repressed splicing similar to the wild-type RBM20, this function was mildly but significantly impaired in the D10 mutant (Fig. 5 A). The other reporter system employs RT-PCR-based analysis of mRNAs derived from a transiently transfected exon/intron cassette containing exons 241, 242 and 243 of human titin (Fig. 5 B) (Dauksaite and Gotthardt, 2018). In this system, we also observed that the D10 mutant significantly impaired exon exclusion compared to the wild-type protein (Fig. 5 B).

**Figure 5.**
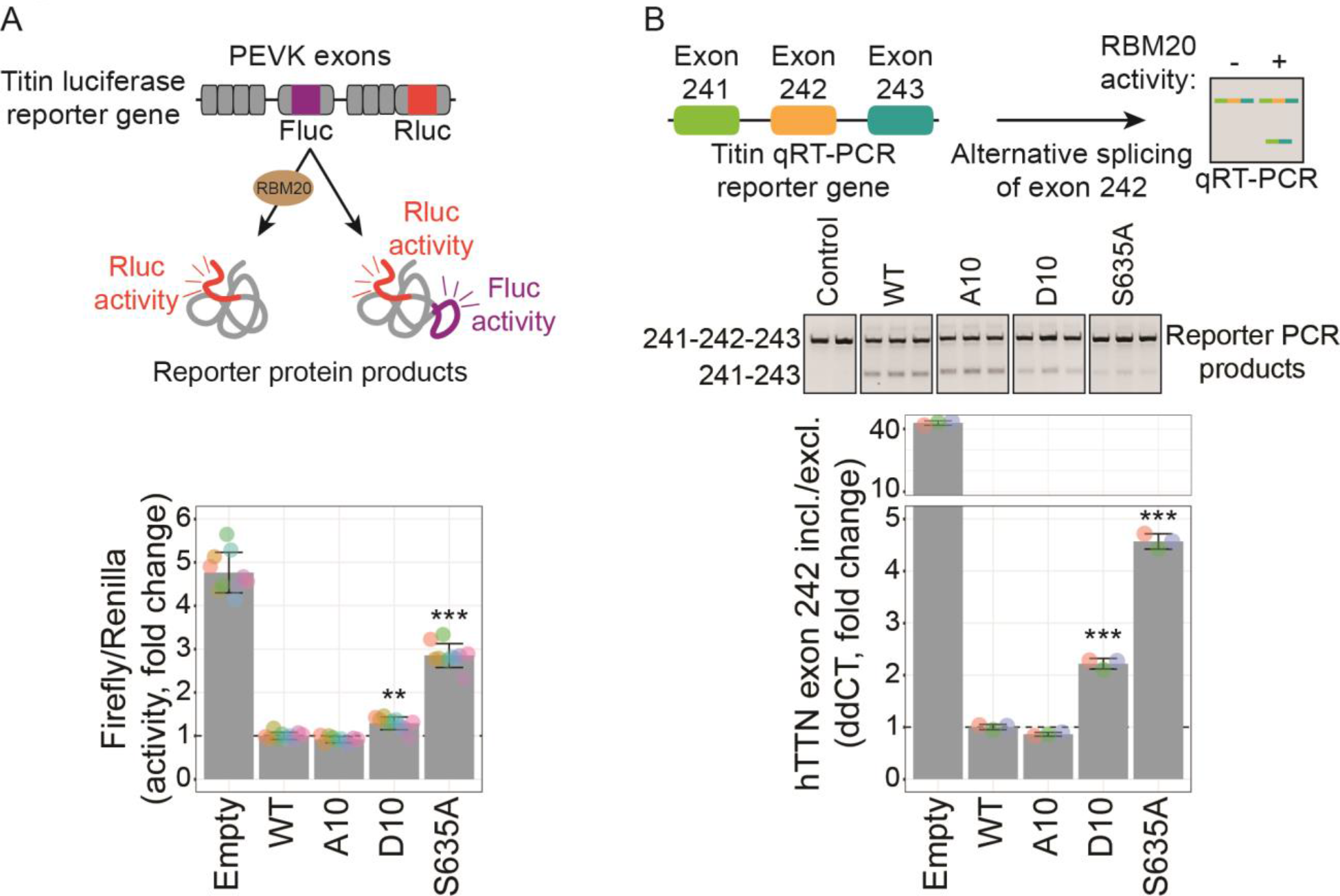
Phosphorylation suppresses RBM20 splicing activity on titin splicing reporters. (A and B) Luciferase- and qRT-PCR-based splicing reporter assay for RBM20 activity on Titin exon inclusion/exclusion. RBM20-regulated Firefly (Fluc) luciferase activity relative to Renilla (Rluc) luciferase activity was quantified in A (n = 10). Inclusion and exclusion of exon 242 was quantified with qRT-PCR in B (n = 3). Data is normalized to the wild-type RBM20 level. Transfection with an empty plasmid was used as control. Data is expressed as the mean (bars) of biological replicates (colored dots) ± SD. Groups were compared by one-way ANOVA and Tukey post-test was used. P-values were considered statistically significant as follow: *p< 0.05; **p < 0.01; ***p< 0.001. P-value significance for the comparisons against the WT group are shown (except for the empty vector control).

In summary, both reporter systems consistently show that the splicing regulatory function of the phosphomimetic D10 mutant is significantly impaired while the non-phosphorylatable A10 mutant is similar to wild-type RBM20. In agreement with BioID and microscopic data, these data indicate that the putative regulatory phosphorylation sites identified by qRIC indeed affects the splicing regulatory function of RBM20.

## Discussion

While mRBPs are central players of posttranscriptional gene regulation, we know relatively little about how their activity is controlled by cellular signaling events. Here, we developed and employed qRIC to study the relationship between phosphorylation of mRBPs and their interaction with mRNA. The idea behind qRIC is to quantify the fraction of the total cellular pool of mRBPs isolated with mRNA, that is, their pull-down efficiency. Comparing this pull-down efficiency of individual phosphorylation sites with the pull-down efficiency of the host protein can then reveal sites with regulatory potential. We show that qRIC data faithfully recapitulates known regulatory phosphorylation sites and identifies novel sites in RBM20 that affect splicing function.

A key advantage of qRIC is that it enables systematic prioritization of phosphorylation sites in mRBPs that are likely to have regulatory function. The methodology therefore complements phosphoproteomic studies that routinely quantify changes in thousands of phosphorylation sites but do not provide direct insights into their possible function. In fact, it is now thought that many phosphorylation sites may not be functionally relevant, highlighting the need for scalable methods to identify functional sites (Ochoa *et al*., 2020). qRIC is conceptually similar to hotspot thermal profiling in the sense that in both cases the fraction of the total pool of proteins is relatively quantified (Huang *et al*., 2019; Smith *et al*., 2021). However, in contrast to assessing protein thermal stability, qRIC assesses the impact of modifications on mRNA-binding and is thus a read-out for sites involved in posttranscriptional regulatory processes.

Despite its strengths, it is also important to keep the limitations of the methodology in mind. Most importantly, qRIC only reveals correlations between modification state and RNA-binding and does not establish causal relationships. Therefore, while it is tempting to assume that phosphorylation of sites with positive (or negative) log2 delta efficiencies causes increased (or decreased) RNA-binding, this is not the only plausible interpretation. It is also possible that changes in RNA-binding cause differences in the phosphorylation state, for example by exposing a site to kinases or phosphatases. Also, correlations between phosphorylation and RNA-binding can be indirect. For example, the phosphorylation of S202 in ELAVL1 we observed to correlate with increased mRNA-binding (Fig. 3) actually causes nuclear retention of the protein and thereby decreases mRNA-binding in the cytosol indirectly (H. H. Kim *et al*., 2008; Grammatikakis, Abdelmohsen and Gorospe, 2017). It is also salient to bear in mind that pull-down efficiencies are affected by factors such as UV cross-linking and mRNA isolation with oligo(dT) beads and therefore do not provide a direct read-out for mRBP-mRNA interaction (Vieira-Vieira and Selbach, 2021). Hence, while qRIC efficiently prioritizes modification sites with regulatory potential, establishing causal relationships still requires follow-up experiments.

Another limitation of our implementation of qRIC is that the peptides UV cross-linked to RNA are systematically missed. In future experiments, chemical cleavage of RBP-RNA cross-links could be used to circumvent this bias (Bae *et al*., 2020). Moreover, instead of oligo(dT) enrichment, future implementations of qRIC could employ organic phase separation methods to broaden the scope of qRIC from mRNAs to other classes of cellular RNAs (Urdaneta *et al*., 2019; Smith *et al*, 2020). Also, combining qRIC with analysis of other types of PTMs (methylation, acetylation, ubiquitination, SUMOylation etc.) could be used to assess their impact on RNA-binding. Finally, implementing qRIC with TMT labeling and deep fractionation might improve coverage of modified sites (Hogrebe *et al*., 2018).

Our data revealed an unexpected impact of phosphorylation on the function of the splicing factor RBM20. This protein has emerged as both a key regulator of cardiac splicing and an important disease protein in dilated cardiomyopathy (Guo *et al*., 2012). Multiple phosphorylation sites in RBM20 have been identified, but previous studies mainly focused on phosphorylation of serine residues in the RS region of the protein (Murayama *et al*., 2018; Sun *et al*., 2020). This region is of special interest since it harbours most known pathogenic RBM20 mutations (Lennermann, Backs and van den Hoogenhof, 2020). Here, we identified several sites with regulatory potential in the disordered C-terminal half of RBM20 outside the RS region. In contrast to the presumed function of phosphorylation in the RS region, introducing phosphomimetic mutations at these more C-terminal sites shifts the subcellular distribution of RBM20 to the cytosol (Fig. 4). This highlights the important fact that PTM functions are typically site-specific (Krug *et al*., 2019). Interestingly, we find that both the phosphomimetic D10 RBM20 mutant and the disease-causing S635A mutant show increased interactions with cytosolic stress granules (Fig. 4). However, while the S635A mutant shows markedly reduced interaction with splicing factors and is almost exclusively cytosolic, the D10 mutant still associates with splicing factors, is partially nuclear and has a less severe impact on splicing. This indicates that the subcellular distribution of RBM20 does not only depend on the RS domain. In fact, a previous study already reported that the RS region is not strictly required for nuclear localisation of RBM20 (Filippello *et al*., 2013).

It is tempting to speculate that phosphorylation at the regulatory sites identified here fine-tunes RBM20 function. This could be physiologically relevant since cardiac splicing is known to change dynamically, for example during development (Lahmers *et al*., 2004). Also, it would be interesting to identify the kinase(s) and phosphatase(s) involved. It might be possible that inhibiting or reducing RBM20 phosphorylation shifts pathogenic variants from the cytosol towards the nucleus, which could restore splicing function. Corresponding kinase inhibitors might therefore even have therapeutic potential in dilated cardiomyopathy patients. One important limitation of our data is that the phosphomimetic D10 mutant is not biochemically identical to WT RBM20 phosphorylated at the corresponding sites. While it is reassuring that the A10 mutant does not seem to be functionally impaired (Fig. 4 and 5), this does not rule out issues arising from the artificial nature of phosphomimetic mutations (Dephoure *et al*., 2013). It would also be interesting to study the impact of RBM20 phosphorylation on its function in a more physiological model system such as cardiomyocytes (Guo *et al*., 2012).

## Acknowledgements

We would like to thank Martha Hergeselle, Janine Fröhlich and Christian Sommer for excellent technical assistance, James Knight for providing the proximity map data, Dushyandi Rajendran and Anne-Claude Gingras for providing the plasmids (all at Lunenfeld-Tanenbaum Research Institute, Toronto, Ontario Canada). We also thank Markus Landthaler (MDC, Berlin Germany) and lab members for helpful discussions.

## Author contributions

Conceptualization, C.H.V-V. and M.S.; Methodology, C.H.V-V. and M.S.; Formal Analysis, C.H.V-V.; Investigation, C.H.V-V. and V.D.; Resources, M.G. and M.S.; Writing – Original Draft, C.H.V-V. and M.S.; Writing – Review & Editing, C.H.V-V., V.D., M.G., and M.S.; Visualization, C.H.V-V.; Supervision, M.G. and M.S.; Project Administration, M.S.; Funding Acquisition, M.S.

## Inclusion and diversity statement

One or more of the authors of this paper self-identifies as an underrepresented ethnic minority in science.

## STAR*METHODS

### RESOURCE AVAILABILITY

#### Lead contact

Further information and requests for resources and reagents should be directed to and will be fulfilled by the lead contact, Matthias Selbach.

#### Materials availability

Plasmids, cell lines and other unique/stable reagents generated in this study are available from the lead contact without restriction.

#### Data and code availability

● Proteomic raw datasets have been deposited to the ProteomeXchange Consortium via the PRIDE partner repository and are publicly available as of the date of publication. Accession numbers are listed in the key resources table. Microscopy data reported in this paper will be shared by the lead contact upon request.
● This paper does not report original code.
● Any additional information required to reanalyze the data reported in this paper is available from the lead contact upon request.

#### Experimental model and subject details

##### Cell lines

All cell lines created in this study are Flp-In T-REx 293 (Thermo Fisher Scientific, cat# R78007) derivatives and listed in the key resources table. All cell cultures were periodically checked for the presence of Mycoplasma by PCR. Commercial cell lines were obtained directly from the commercial sources and have therefore not been authenticated in-house. HEK293T and Flp-In T-REx 293 derivative cell lines were cultured in high glucose Dulbecco’s modified Eagle’s medium (DMEM) complemented with glutamax (Thermo Fisher Scientific, cat# 10569010) and 10% fetal bovine serum (FBS) (PAN-Biotech, cat# P30-3306). HEK293 cells were maintained in high glucose DMEM supplemented with 10 % (v/v) FBS (Sigma-Aldrich, cat# F7524), 100 U/ml penicillin and 100 µg/ml streptomycin. Cells were cultivated at 37 °C in the presence of 5 % CO2 in a humidified, dark chamber. For SILAC labeling, cells were grown in arginine- and lysine-free DMEM (PAN-Biotech, cat# P04-02505) containing 10 % (v/v) dialyzed FBS (PAN-Biotech, cat# P30-2102) and 1 % glutamax (Thermo Fisher Scientific, cat# 35050038) in the presence of 0.2 and 0.8 mM of L-arginine (Arg0) (Sigma-Aldrich, cat# A6969) and L-lysine (Lys0) (Sigma-Aldrich, cat# L8662), L-[13C6]-arginine (Arg6) (Sigma-Aldrich, cat# 643440) and L-[2H4]-lysine (Lys4) (Cambridge Isotope Laboratories, cat# DLM-2640-PK) or L-[13C6,15N4]-arginine (Arg10) (Sigma-Aldrich, cat# 608033) and L-[13C6,15N2]-lysine (Lys8) (Silantes, cat# 211604102), respectively. To achieve complete incorporation of SILAC amino acids, cells were cultured in SILAC medium by continuously passing and growing cells at low confluency (<50%) for at least eight days corresponding to roughly seven doubling. Labeling efficiency was confirmed by mass spectrometry.

#### Method Details

##### qRIC and sample preparation for (phospho)proteome analysis

Fully labeled SILAC Human Embryonic Kidney 293T (HEK293T) cells were grown in seven 15-cm dishes per label until 60 % confluency. Cells were washed on the plate with 5 mL of ice-cold PBS and irradiated with 0.15 J/cm² of 254 nm UV light for *in vivo* cross-linking of RNPs. After cross-linking, cells were collected in 50 mL of ice-cold dPBS by scrapping the plates. Light-labeled cells were kept as input material and heavy-labeled cells were used for RIC. After removal of supernatant, cell pellets were resuspended in 7 mL of lysis buffer (100 mM Tris pH 7.5, 500 mM LiCl, 1 % LiDS, 10 mM EDTA pH 8.0) with freshly added 5 mM of DTT, protease inhibitors (1 pill for every 50 mL final volume) (Roche, cat# 4693159001) and 1:100 dilution of phosphatase inhibitor cocktails 2 and 3 (Sigma-Aldrich, cat# P5726 and P0044). Samples were lysed and homogenized by passing it ten times through a 21-gauge needle and five times through a 26-gauge needle. For the experiment where a non-crosslinking control was included, half the number of medium-heavy labeled cells were similarly grown and the lysate was mixed with the heavy labeled lysate before continuing with RIC. A reverse label experiment was simultaneously performed by swapping the light and heavy SILAC samples for pull-down and input sample. In total, three forward and three reverse experiments were performed, but only one included the medium-heavy labeled non-crosslinking control.

For the pull-down of mRNA and cross-linked mRBPs, 15 mL of oligo(dT) magnetic beads (New England Biolabs, cat# S1419S) (0.5 mL per 0.1 g of cell pellet) were pre-washed with 30 mL of lysis buffer before incubation with cell lysates for 3 hours at room temperature. Beads were collected with magnetic racks and washed three times with 7 mL of lysis buffer and two times with 5 mL of NP-40 washing solution (50 mM Tris pH 7.5, 140 mM LiCL, 2 mM EDTA pH 8.0, 0.5 % NP-40) with freshly added 5 mM of DTT, protease inhibitors (1 pill for every 50 mL final volume) and 1:100 dilution of phosphatase inhibitor cocktails 2 and 3. Beads were further washed three times with 1 mL of 50 mM ammonium bicarbonate (ABC) solution and resuspended in 600 µL of ABC solution.

In parallel, input samples were resuspended in 100 mM Tris pH 7.5 with 0.5 % SDS and freshly added protease inhibitors (1 pill for every 50 mL final volume) and 1:100 dilution of phosphatase inhibitor cocktails 2 and 3. Input samples were lysed by boiling at 98 °C for 5 min. Excessive DNA and RNA were removed by letting the sample cool down and digestion with 1 µL of Benzonase (Merck, cat# 101695) at 37 °C for 30 min. Proteins were precipitated by addition of nine volumes of absolute ethanol and overnight incubation at -20 °C followed by 30 min centrifugation at 20000 g at 4 °C. Input protein pellets were resuspended in 200 µL of 2 M Urea, 6M Thiourea solution with freshly added 10 mM of DTT for 5 min at room temperature. The light labeled input protein sample (corresponding to 1 % of the initial input) was mixed with the oligo(dT) magnetic beads used in RIC of heavy labeled cells. Proteins were alkylated in the dark with 55 mM of iodoacetamide or chloroacetamide for 20 min at 25 °C. For lysis, proteins were incubated with 10 µg of lysyl endopeptidase (Wako Chemicals, cat# 129-02541) at 25 °C for 2 hours and incubated with 10 µg of trypsin (Promega, cat# V5113) under constant agitation at 25 °C for 16 hours in the dark. Peptides were acidified with 1% (v/v) trifluoroacetic acid and desalted with C18 Stage Tips (Rappsilber, Mann and Ishihama, 2007). A large fraction of the peptide sample (90 %) was used for enrichment of phosphopeptides. Remaining peptides were eluted with 50% acetonitrile 0.1% formic acid, dried and resuspended in 3% acetonitrile, 0.1% formic acid.

For the experiment that included non-crosslinked cells in the medium-heavy channel, peptides were fractionated by cation-exchange chromatography prior to LC-MS/MS analysis to improve coverage. For that, peptides were loaded into a microcolumn tip packed with strong cation exchange Empore discs (3M and cat# 66889) and washed two times with a 0.5 % formic acid, 20 % acetonitrile aqueous solution. Three fractions were collected by eluting peptides with increasing concentration of salt: 125, 250 and 500 nM of ammonium acetate in 0.5 % formic acid, 20 % acetonitrile solution. Peptide solutions were further diluted to final 4 % acetonitrile concentration and pH acidified with addition of formic acid to a 5 % final concentration before desalting with C18 Stage Tips for a second time.

##### Phosphopeptide enrichment in qRIC samples

Desalted peptides were eluted from Stage Tips in 300 µL of loading buffer (80% acetonitrile [vol/vol] and 6% trifluoroacetic acid [vol/vol]). Phosphopeptides were enriched using a microcolumn tip packed with 1 mg of TiO2 Titansphere (GL Sciences, cat# 5020-75010). At 4 °C, the TiO2 column was equilibrated by passing through 20 µL of the loading buffer with centrifugation at 100 g. Sample solution was completely loaded on the TiO2 column via multiple consecutive steps of centrifugation at 100 g. Next, the TiO2 column was washed with 20 µL of the loading buffer, followed by 20 µL of 50 % acetonitrile (vol/vol), 0.1 % trifluoroacetic acid (vol/vol) solution. The bound phosphopeptides were eluted using successive elution with 30 µL of 5 % NH3.H2O solution (fraction 1) followed by 30 µL with 5 % piperidine solution (fraction 2). Each fraction was collected into a fresh tube separately containing 30 µL of 20 % formic acid and further acidified with formic acid until pH smaller than 2 was obtained. The phosphopeptides were desalted with C18 Stage Tips prior to LC-MS/MS analysis.

##### NanoLC-MS/MS analysis of digested peptides from qRIC experiments

For LC-MS/MS analysis, desalted peptides were eluted from Stage Tips with 50 % acetonitrile 0.1 % formic acid solution, dried and resuspended in 3 % acetonitrile 0.1% formic acid. Peptide concentration was determined based on 280 nm UV light absorbance. Reversed-phase liquid chromatography was performed employing an EASY nLC II 1200 (Thermo Fisher Scientific) using self-made 20 cm long C18 microcolumns packed with ReproSil-Pur C18-AQ 1.9-μm resin (Dr. Maisch, cat# r119.aq.0001) connected on-line to the electrospray ion source (Proxeon) of an HF-X Orbitrap mass spectrometer (Thermo Fisher Scientific). The mobile phases consisted of 0.1 % formic acid 5 % acetonitrile solution (Buffer A) and 0.1 % formic acid 80 % acetonitrile solution (Buffer B). Peptides were eluted at a flow rate of 250 nL/min over 44 to 214 min of increasing Buffer B concentration. Settings for data dependent mass spectrometry analysis were as follow: positive polarity, one full scan (resolution, 60000; m/z range, 350-1800; AGC target, 3e6; max injection time, 10 ms) followed by top 20 MS/MS scans using higher-energy collisional dissociation (resolution, 15000; m/z range, 200-2000; AGC target, 1e5; max injection time, 22 ms; isolation width, 1.3 m/z; normalized collision energy, 26). Ions with an unassigned charge state, singly charged ions, and ions with charge state higher than six were rejected. Former target ions selected for MS/MS were dynamically excluded within 20 s.

##### Processing mass spectrometry data with MaxQuant

All raw files from the same experiment were analyzed together with MaxQuant software (v1.6.0.1) (Cox and Mann, 2008) using default parameters. For increasing transparency and reproducibility of data analysis the “mqpar.xml” file generated by MaxQuant was deposited together with the raw data. Briefly, search parameters used for identification and quantification included two missed cleavage sites, cysteine carbamidomethyl as fixed modification, and the following variable modifications: methionine oxidation, protein N-terminal acetylation, and asparagine or glutamine deamidation. Up to three variable modifications per peptide were allowed. Lys0 and Arg0, Lys4 and Arg6, or Lys8 and Arg10 were set as multiplicity labels. Peptide mass tolerance was 20 and 4.5 ppm for first and main search, respectively. Database search was performed with Andromeda embedded in MaxQuant against the UniProt/Swiss-Prot Human proteome (downloaded in January 2019) with common contaminant sequences provided by MaxQuant. False discovery rate was set to 1% at peptide spectrum match and protein levels. Minimum peptide count required for protein quantification was set to two. The “Requantify” option was turned on. An identical MaxQuant search but with the “Requantify” option off was performed by partial reprocessing of search post peptide searches (starting from step “Re-quantification”). The second run (with Requantify off) was used for identification and exclusion of unscrupulous ratios (defined as ratios between two requantified values). Results from both searches with and without Requantify are provided. Mass spectrometry proteomics and phosphoproteomics raw data and MaxQuant output tables for the qRIC experiments have been deposited to ProteomeXchange Consortium (http://proteomecentral.proteomexchange.org) via the PRIDE partner repository with the dataset identifier PXD027137. Similarly, raw data and MaxQuant output tables for the BioID experiment have been deposited with the identifier PXD027138.

##### Analysis of qRIC results and calculation of delta pull-down efficiency

The “proteinGroups.txt”, “peptides.txt” and “Phospho (STY)Sites.txt” tables from a single MaxQuant run including all three experiments were used for data analysis of results. Potential contaminants, reverse database hits, and peptides only identified with modification were excluded. Log2 SILAC ratios from reverse labeled experiments were mathematically inverted to reflect pull-down over input ratios. For analysis, only hits with quantified SILAC ratios in at least one forward and one reverse experiment were kept. The mean of log2 pull-down over input ratios from all three forward and reverse label experiments were taken individually, while the grand mean was taken as the mean of means in log2 space. To calculate pull-down efficiencies, the pull-down over input ratios (linear scale) were multiplied by the percentual amount of spike-in input material relative to the total amount of material used in RIC. Results in all three experiments were highly reproducible overall. As an additional quality filter, we discarded a few irreproducible measurements (difference in the forward and inverted reverse SILAC ratios higher than 4 and 8 fold at protein and phosphopeptide level, respectively). This removed 256 proteins and 63 phosphopeptides in total but only 4 and 16, respectively, above the 1 % pull-down efficiency threshold for delta efficiency calculation.

Protein and phosphosite information was merged in a single table and the delta pull-down efficiency in each experiment was calculated by subtracting the pull-down over input log2 ratio of individual phosphosite and its host protein. As a result, in log2 scale, positive (or negative) delta pull-down efficiency indicates that the phosphopeptide was pulled down more (or less) efficiently than the host protein. As before, mean delta efficiency in each experiment label was calculated and the grand mean taken. In most figures, the grand mean is plotted. Hits were deemed specific when the mean delta efficiency was above 2 fold change in either direction.

##### Sensitivity and specificity analysis of qRIC

Proteins quantified in qRIC were compared to the set of annotated mRBPs (named “all mRBPs”) obtained from Hentze et al. (2018). The subset from these that has been identified in HEK293 cells were also selected for comparison (named “HEK mRBPs”). In addition, by uniting the set of mRBPs with the set of manually curated RBPs in Gerstberger et al (2014) we generated a full list of all RBPs (named “all RBPs”). For comparison with my qRIC data in Figure 1, we matched datasets by gene names. In Figure 1E, receiver operating characteristic analysis was performed with the R package pROC (Robin *et al*., 2011) using annotated RBPs as positives and non-annotated proteins as negatives.

##### Analysis of pull-down efficiencies correlation with specific RBP features

Protein domain annotations were obtained from the Pfam database downloaded in May 2016 (Mistry *et al*., 2021b) for all proteins quantified in qRIC. Among all domains, RBDs were selected based on the manually curated list provided in Gerstberger et al (2014). For comparison with qRIC data in Figure 2C and 2D, we matched proteins in both datasets by Uniprot ID. For the comparison of pull-down efficiencies of RBPs in our data with corresponding CLIP-seq data in Figure 2 E and S2 A, the entire POSTAR2 database of uniformly analysed PAR-CLIP experiments was downloaded in August 2019 (Zhu *et al*., 2019). Experiments performed in HEK293 and HEK293T were not differentiated. Other types of CLIP-seq data available in the POSTAR2 database were not used for the analysis as results for only few proteins exist. Similarly, the entire dataset of eCLIP experiments was downloaded from ENCODE (Davis *et al*., 2018) in January 2021 and used in Figure S2 B and C. Peaks in all datasets not mapped to human chromosomes (chromosomes 1 to 21, X, Y and mitochondrial) were excluded from analysis. CLIP-seq and qRIC datasets were merged by gene names.

##### Generation of HEK293 Stable Cell Lines Expressing mutant RBM20 variants

For generation of plasmids for expression of RBM20 C-terminally fused to the BirA* (a.k.a. BioID) and FLAG tag, the wild-type RBM20 coding sequence and that of the S635A variant without the stop codon have been previously cloned Maatz et al (2014). D10 and A10 mutant sequences were synthesized by BioCat based on the wild-type sequence by altering the corresponding serine codons to aspartic acid or alanine codons, respectively. Sequences were validated by sequencing. Following manufacturer’s instruction, the synthetic sequences were cloned into the pDONR221 using Gateway Clonase II system (Thermo Fisher Scientific) and transferred to pDEST_pcDNA5_BirA-FLAG_Cterm (Couzens *et al*., 2013) for protein expression. The resulting vectors (pEXPR_RBM20-WT_C-term_BirA_FLAG, pEXPR_RBM20-D10_C-term_BirA_FLAG, pEXPR_RBM20-A10_C-term_BirA_FLAG, pEXPR_RBM20-S635A_C-term_BirA_FLAG) were used to generate stable HEK293 Flp-In T-Rex cells lines overexpressing C-terminally BioID- and FLAG-tagged WT RBM20, or A10, D10 and S635A RBM20 variants. For that, HEK293 Flp-In T-Rex cells were transfected in a 6-well format by mixing 200 µL of FBS-free medium with 1.5 µg of total plasmid DNA (2:1 ratio of pOG44 to destination vector) and 3.75 µL of 40 kDa linear polyethylenimine (PEI40) (Polysciences, cat# 24765-2). After a 15 min incubation, the transfection mixture was added to the cells. Cells were re-seeded into 10 cm dishes after 48 hours and allowed to attach overnight. Hygromycin (200 µg/mL) was added the next day and the cells were selected for 18 days by the addition of fresh hygromycin-containing cell culture media every 2-3 days resulting in expansion of monoclonal colonies. Monoclonal colonies were carefully transferred by pipetting cells into 6 cm dishes for expansion in media containing Hygromycin. Cells were confluent after 12 days and cultivated for further experimentation in media without the addition of Hygromycin. Expression of the mutant proteins was validated after treatment of cells for 24 hours with 1 µg/mL of Tetracycline by immunodetection of the FLAG tag in the expected product size in an electrophoretic gel, and detection of specific peptides via shotgun proteomics when possible.

##### Proximity interactors investigation with BioID

Following the experimental design in Figure 4 B, SILAC labeled cells expressing RBM20 variants were grown in two 15-cm dishes to 25% confluency per experimental group and incubated for 24 hours with 1 µg/mL tetracycline to induce expression of RBM20 variants. Light-labeled WT-expressing cells were used as control in the experiment 1 (see Figure 4 B) by omitting tetracycline from the media and served as an uninduced control for background binding. The relative SILAC quantification allowed for comparison of proteins that have been proximity labeled by the transiently expressed constructs. After the induction period all cell lines were incubated for 24 hours in the cell culture medium containing biotin. Then, cells were lysed in lysis buffer (50 mM Tris-HCl pH 7.5, 150 mM NaCl, 1% Triton X-100, 1 mM EDTA, 1 mM EGTA, 0.1% SDS) with freshly added protease inhibitors (1 pill for every 10 mL final volume) and 0.5 % sodium deoxycholate. Excessive DNA and RNA were removed by digestion with 2 µL of Benzonase at 37 °C for 20 min. A small fraction of the cleared input lysate was precipitated by addition of nine volumes of absolute ethanol and overnight incubation at -20 °C followed by 30 min centrifugation at 20000 g at 4 °C. Input protein pellets were resuspended in 2 M Urea, 6 M Thiourea solution with freshly added 10 mM of DTT, alkylated in 55 mM iodoacetamide, digested with LysC for 3h at 25°C, diluted four times with 25 mM ammonium bicarbonate buffer and digested overnight with Trypsin at 25°C. Remaining input lysate was used for enrichment of biotinylated proteins by incubation with streptavidin-sepharose beads (Sigma-Aldrich, cat# GE17-5113-01) for 3 hours at 4 °C. Beads were washed once with lysis buffer, twice with washing buffer (50 mM HEPES-KOH pH 8.0, 100 mM KCl, 10% glycerol, 2 mM EDTA, 0.1 % NP-40) and six times with 25 mM ammonium bicarbonate buffer to completely remove detergents from the sample. Beads were eluted in ammonium bicarbonate and proteins were digested with trypsin. Beads were removed by centrifugation and peptides were acidified with 1% (v/v) trifluoroacetic acid and desalted with C18 Stage Tips.

Mass spectrometry analysis was performed similarly as before (see “NanoLC-MS/MS analysis of digested peptides from qRIC experiments”). The main difference was that peptides were analysed in a Orbitrap Exploris 480 mass spectrometer (Thermo Fisher Scientific) in Peptide mode and with an associated FAIMS device (Thermo Fisher Scientific) installed for pseudo ion mobility peptide separation. Peptides elution gradient was 110 min long. Settings for data dependent mass spectrometry analysis were as follow: positive polarity, one full scan (resolution, 120000; m/z range, 350-1800; normalized AGC target, 300 %; max injection time, 30 ms) followed by top 20 MS/MS scans using higher-energy collisional dissociation (resolution, 7500; m/z range, 200-2000; normalized AGC target, 100 %; max injection time, 25 ms; isolation width, 1.3 m/z; normalized collision energy, 28). Three successive rounds of a full scan followed by 20 MS/MS scans were constantly performed throughout the scanning phase with rotating correction voltages of -40, -60 or -80 V. Scans from single correction FAIMS voltages were collected into MzXML files using the FAIMS MzXML Generator tool provided by the Coon lab: https://github.com/coongroup/FAIMS-MzXML-Generator.

The “proteinGroups.txt” from a single MaxQuant run including all MzXML files were used for data analysis of results with MaxQuant (see *Processing mass spectrometry data with MaxQuant*). Potential contaminants, reverse database hits, and proteins only identified by modified peptides were excluded. Log2 SILAC ratios from reverse labeled experiments were mathematically inverted to reflect the induced over uninduced or the mutant over WT experiments. For analysis, only hits with quantified SILAC ratios in both forward and reverse experiments were considered. Proteins with fold change higher than 2 in both forward and reverse experiments were considered significant when comparing induced and uninduced WT cell lines. To correct for proteome differences in cell lines expressing the RBM20 variants, biotin-enriched samples were normalized by subtracting log2 fold changes and the respective log2 fold change in the input proteome sample. Proteins with proteome-normalized fold changes higher than 1.5 in both forward and reverse experiments comparing mutant and WT variants were considered significant. Data presented in Fig. 4 E and F are the mean values between forward and reverse experiments. Gene ontology enrichment of cellular components, molecular functions and biological processes was performed with Metascape (Zhou *et al*., 2019).

##### RBM20 immunostaining and imaging

Cells stably expressing RBM20 variants were seeded on coverslips coated with poly-L-lysine. Variant protein expression was induced in media containing 1 µg/mL tetracycline for 24 hours before cells were fixed for 15 min with 4 % paraformaldehyde at room temperature. Fixed cells were permeabilized for 10 min with 0.5 % Triton in PBS at room temperature and nonspecific protein binding was blocked by incubation in 1.5 % BSA PBS solution for 1 hours with low shaking. Cells were immuno stained by incubation for 1 hour at room temperature with a specific antibody against FLAG (1:1000 dilution) conjugated to Alexa 488 (Cell Signaling, cat# 5407S). Nucleus was stained with DAPI (Sigma-Aldrich, cat# D9564). Images were acquired by Leica DM5000b microscope with an HCX PL FL 20x/0.50 objective. Images were further processed with Fiji ImageJ (Schindelin *et al*., 2012). For quantification of cells with nuclear, cytosolic or widespread RBM20 subcellular localization at least 250 cells were counted in each group.

##### Cell-based Luciferase TTN splicing reporter assay

For transfection, HEK293 cells were seeded on 96-well plates and transfected with a total of 200 ng of plasmid DNA of which 1 ng was splice reporter PEVK Ex4-13 (Guo *et al*., 2012) plus a corresponding amount of plasmid for expression of RBM20 variants (see “Generation of HEK293 Stable Cell Lines Expressing mutant RBM20 variants”) or control plasmid (pcDNA3.1) in a 20x molar excess. To deliver plasmid DNA, we used PEI40 at a 1:3 ratio (DNA: PEI40). Plasmids and PEI40 in FBS-free medium were incubated for 15 min before the transfection mixture was added to the cells. Cells transfected were at a confluence of 50-60%. Each transfection experiment was repeated ten times and cell viability was measured 60 hours post-transfection using PrestoBlue (Thermo Fisher Scientific, cat# A13261). Luciferase activity was measured 60 hours post-transfection using the Dual-Luciferase® Reporter Assay System (Promega) on an Infinite® M200 Pro (TECAN) plate reader. Ratios of firefly to renilla luciferase activity were normalized to the WT RBM20 expressing cells. All data are expressed as the mean of biological replicates (n = 10) ± SEM. Group comparisons were analyzed by one-way ANOVA and Bonferroni post test. P values were considered statistically significant as follows: *P < 0.05; **P < 0.01; ***P < 0.001.

##### Cell-based qRT-PCR TTN splicing reporter assay

For transfection, HEK293 cells were seeded on 6-well plates and transfected with PEI40 at a 1:3 ratio (DNA:PEI40). Plasmids for the TTN^241-3^ splicing reporter system (Dauksaite and Gotthardt, 2018) and for expression of RBM20 variants (see “Generation of HEK293 Stable Cell Lines Expressing mutant RBM20 variants”) were mixed with PEI40 in FBS-free medium and incubated for 15 min before the transfection mixture was added to the cells. Each transfection experiment was performed using three technical replicates and repeated three times. Transfected cells were equally divided for RNA and protein analysis 48 hours after transfection. Similar levels of expression for all RBM20 variants was validated by immunodetection of RBM20 using a specific antibody (Abcam, cat# ab233147) (data not shown). RNA was isolated from cells using the TRIzol reagent following instructions from the manufacturer (Thermo Fisher Scientific, cat# 15596026). Preparations with less than 2 μg of total RNA were treated with DNase I (Thermo Fisher Scientific, cat# EN0521) and first-strand cDNA was synthesized using High-Capacity RNA-to-cDNA kit (Applied Biosystems, cat# 4387406). Quantitative RT-PCR was performed using SYBR Green master mix (Applied Biosystems, cat# 4309155) in a 7900 HT cycler (Applied Biosystems). qRT-PCR primers are listed somewhere else (Dauksaite and Gotthardt, 2018). The quantification of the gene expression was performed using the ΔΔCT method. Relative levels of splice isoforms are presented as a ratio of mRNAs, with exon 242 included, versus mRNAs, with exon 242 excluded. The fold change in inclusion/exclusion ratio was obtained when compared to the wild type RBM20. The mean of technical replicate values was used for quantification analysis. All data are expressed as the mean of biological replicates (n = 3) ± SEM. Group comparisons were analyzed by one-way ANOVA and Bonferroni post test. P values were considered statistically significant as follows: *P < 0.05; **P < 0.01; ***P < 0.001.

#### Quantification and statistical analysis

The type of statistical test (e.g., Wilcoxon rank-sum, t-test or ANOVA) and post-hoc p-value correction are annotated in the Figure legends and/or in the Methods and Resources segment specific to the analysis. In addition, statistical parameters such as the value of n, mean/median, SEM, SD and significance level are reported in the Figures and/or in the Figure Legends. Statistical analyses were performed using R Studio as described in Methods and Resources for each individual analysis.

## Supplemental Information

**Figure S1.**
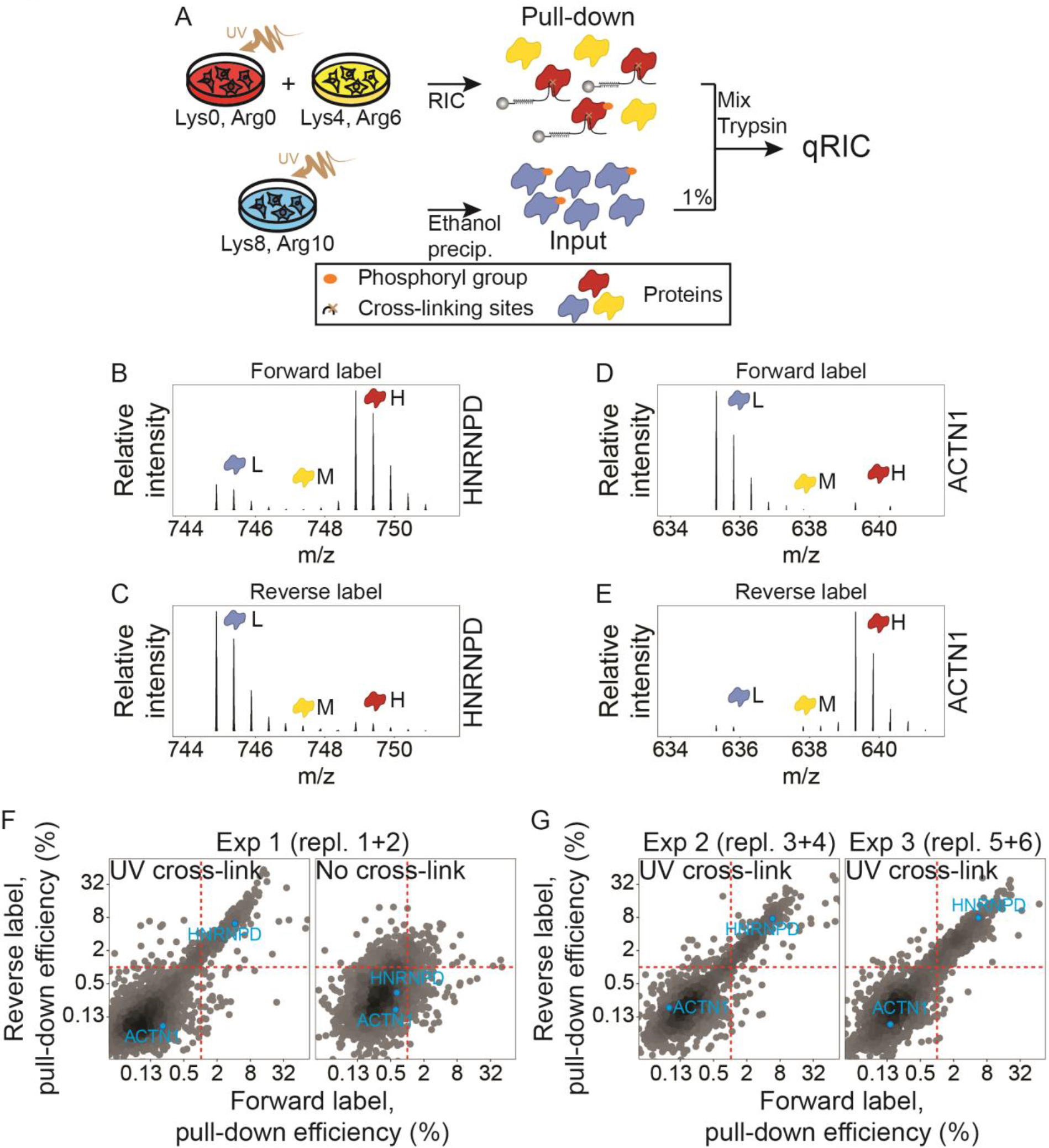
Specific enrichment of previously annotated RBPs. (A) Experimental design of qRIC experiment 1 including the non-crosslinked, medium-heavy SILAC labeled cells. (B, C, D and E) Exemplary spectra for a known RBP (HNRNPD) and a non-RBD (ACTN1). SILAC light (L), medium-heavy (M) and heavy (H) labeled chromatogram peaks are indicated. (F and G) Reproducibility overview of all three qRIC experiments. Pull-down efficiency threshold is indicated by red dashed lines. HNRNPD and ACTN1 are also indicated in the plot.

**Figure S2.**
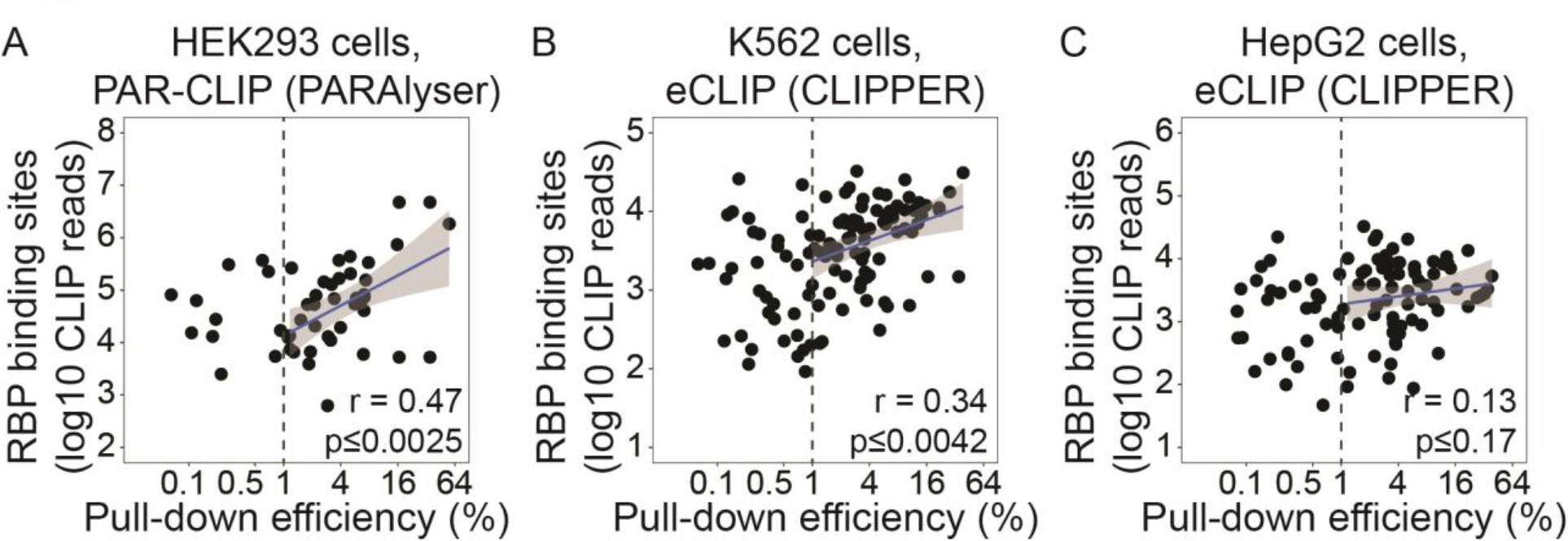
CLIP-seq data correlates with pull-down efficiency in different datasets. (A, B and C) Same as Figure 2 E. Cell line (HEK293, K562 or HepG2), CLIP-seq technology (PAR-CLIP or eCLIP) and peak calling tool (PARAlyser or CLIPPER) are indicated.

**Figure S3.**
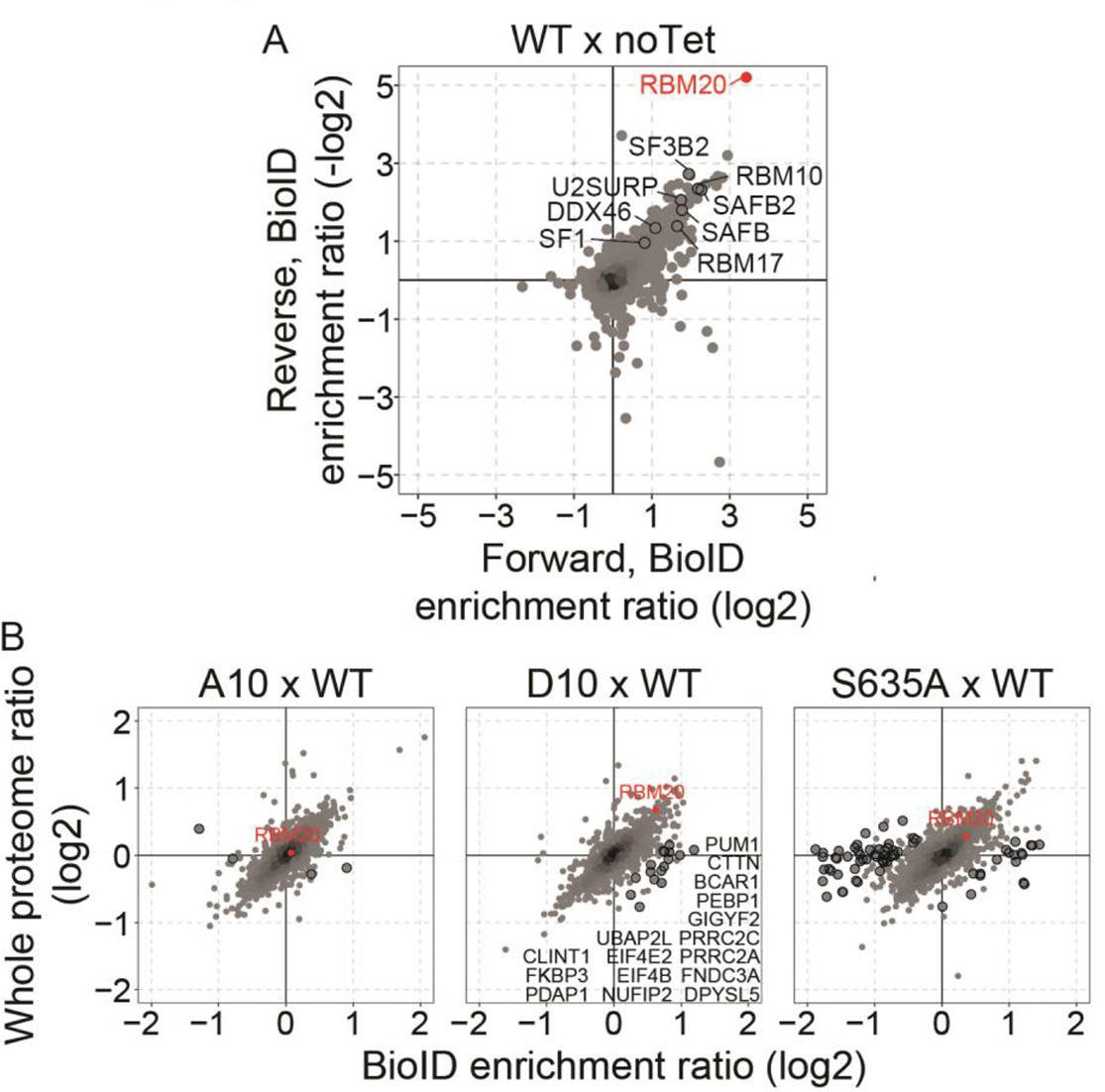
Visualization of BioID data. (A) Reproducibility overview of the BioID enrichment comparing cells expressing the wild-type (WT) RBM20 and non-induced cells (noTet). RBM20 and several splicing factors are highlighted. (B) Overview of biotin enrichment and proteome changes in BioID experiments comparing mutant and wild-type RBM20. RBM20 and proteins preferentially enriched in mutant (over wild-type) or wild-type (over mutant) are highlighted.

Tables legends

Table 1 - Proteins pull-down efficiency in qRIC

Table 2 - Phosphorylation sites pull-down efficiency in qRIC

Table 3 - Delta pull-down efficiency of individual phosphorylation sites

